# Deciphering structural asymmetry of the habenula in the human brain

**DOI:** 10.1101/2022.07.26.501516

**Authors:** Yilamujiang Abuduaini, Yi Pu, Paul M. Thompson, Xiang-Zhen Kong

## Abstract

Functional laterality of the habenula has been suggested in both animal models and the humans. Understanding this evolutionarily conserved brain feature is of fundamental importance and has been attracting attention due to its potential role in human cognition and a variety of neuropsychiatric disorders such as depression and schizophrenia. Deciphering structural asymmetry of the human habenula remains to be challenging. Here, we present a large-scale meta-analysis of the left-right differences in the habenular volume in the human brain with 52 datasets (*N* = 1,427), and also assessed the potential moderating effects of the sampling variability and other methodological factors. Results showed significant heterogeneity in the left-right differences across the datasets, which seems to be mainly due to different MRI scanners and segmentation approaches used. While little evidence was found for the volume asymmetry across all the datasets, the most pronounced and significant leftward asymmetry was found in the datasets from 3 T scanners and when using manual segmentation approaches. We did not find significant disorder-related differences relative to healthy controls in either the left-right asymmetry or the unilateral volume. This study not only provides useful data for future studies of brain imaging and methodological developments related to precision habenula measurements, but also helps to understand potential roles of habenular laterality in health and disorders.

## Introduction

Understanding the hemispheric specialization of the human brain is one of the long-standing questions in human brain research (Duboc et al., 2015; Kong et al., 2022; Toga & Thompson, 2003). Altered asymmetry has been suggested as a potential biomarker in various cognitive and neuropsychiatric disorders, including dyslexia, Alzheimer’s disease, autism, and obsessive-compulsive disorder (Eyler et al., 2012; Kim et al., 2012; Leonard & Eckert, 2008; Menzies et al., 2008). Only recently, large-scale neuroimaging studies have been conducted which aimed to provide a definitive and normative reference of cortical and subcortical asymmetry in healthy and diseased populations (Kong et al., 2022; Ocklenburg et al., 2021). Thus far, existing large-scale studies either focused on mapping of the cortex or relatively large subcortical structures (e.g., hippocampus and thalamus), and the results suggest that such structural asymmetries could explain only a fraction of the variability in functional lateralization (e.g., Dorsaint-Pierre et al., 2006; Guadalupe et al., 2014; Kong et al., 2021). Yet little is known about the nature of the asymmetry for smaller subcortical structures such as the habenula (whose volume is approximately 15-30 mm^3^ in each hemisphere).

The habenula is a bilateral nuclear complex located on the dorsal tip of the thalamus, which is phylogenetically conserved from fish to human (Amo et al., 2010). It has attracted much interest due to its potential role in the evolution of the brain and in its functions. This structure is considered to be essential for allowing animals to switch responses as needed when environments and motivational stages change (Mizumori et al. 2017). The habenular asymmetry has been repeatedly characterized, and this has also been associated with fear responses, sensory processing and various behaviors in different animals including zebrafish, frogs, and rats (Agetsuma et al., 2010; Dreosti et al., 2014; Wree et al., 1981; Guglielmotti & Fiorino, 1998). However, the functional role and the nature of the asymmetry of the habenula in humans remains largely unexplored (Batalla et al., 2017), in part, because of the extremely small size of the habenula and its low anatomical contrast on *in vivo* human brain imaging (Fig. 1).

**Fig. 1:**
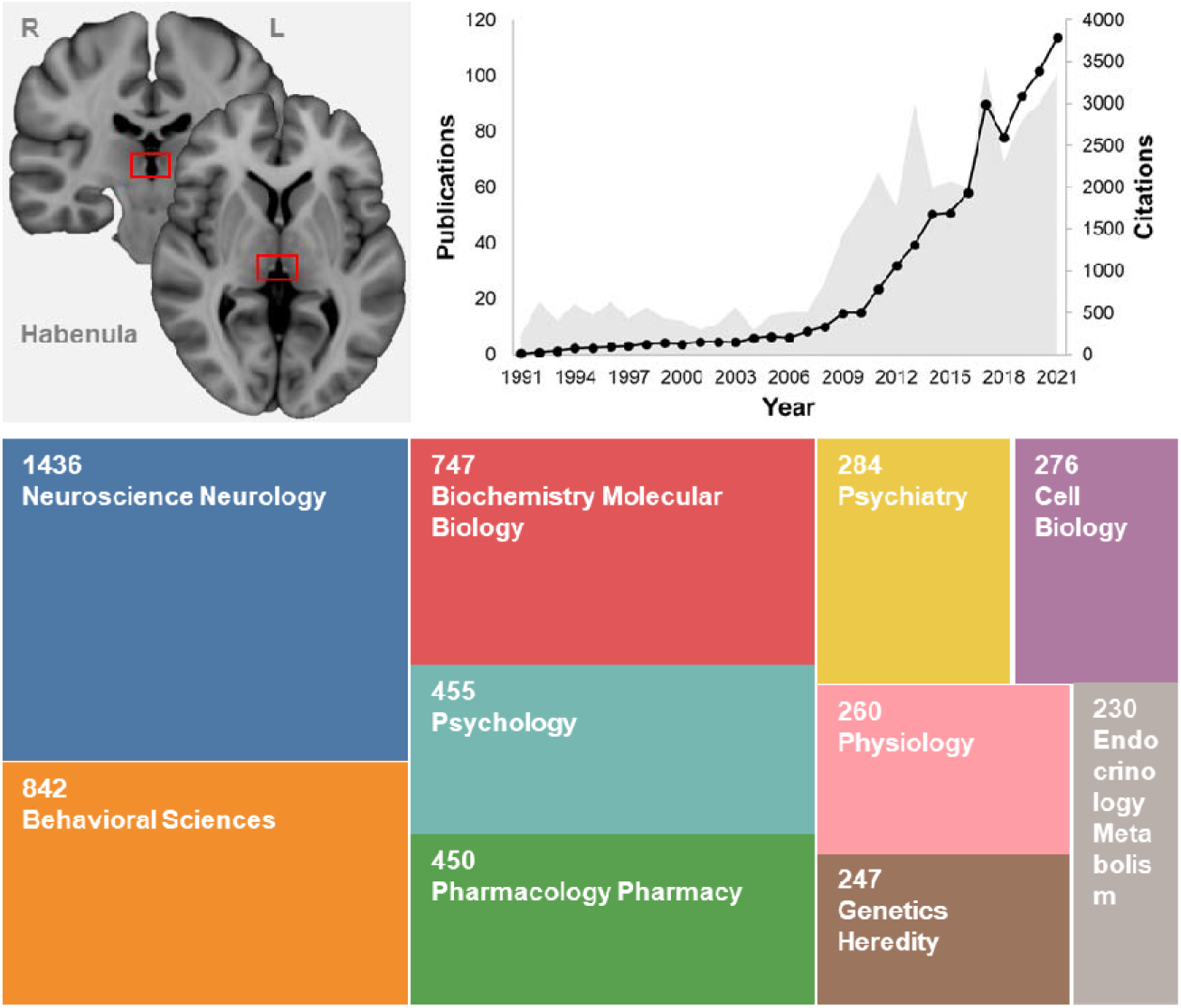
Anatomical location of the habenula in the human brain (top left) and a summary of related studies in the past decades (top right and bottom). Data for the visualization was obtained via Web of Science using search terms “habenula (Title) or habenular (Title)”. The bottom panel indicates the top research areas of the 1550 relevant publications obtained, including Neuroscience/Neurology, Behavioral Sciences, Psychology and Psychiatry. Depression and Schizophrenia are the top brain disorders revealed in the MeSH Headings from Web of Science.

Understanding this evolutionarily conserved brain feature has been attracting attention in different research areas including neuroscience, psychology and psychiatry, due to its potential role in human cognition and a variety of neuropsychiatric disorders such as depression and schizophrenia (see Fig. 1). Some of the leading efforts to understand habenular asymmetry have used high-resolution functional magnetic resonance imaging (fMRI) (Lawson et al., 2013). For example, functional activation in the habenula has been found during the prediction of aversive stimuli (Lawson et al., 2017), which could be, in part, explained by its role in avoidance learning processing (Yoshino et al., 2020). Interestingly, researchers found that the left habenula, compared to the right, seems to be more significantly activated during punishment outcomes, and this activation correlated with higher reward dependence scores and more severe depressive symptoms (Yoshino et al., 2020; Liu et al., 2017). Asymmetric activation of the habenula has also been reported in studies of aversive processing (Lawson et al., 2014; Hennigan et al. 2015). Further evidence in supporting the functional differentiation of the left and right habenula comes from functional connectivity studies. For example, resting-state activity recorded from the left and right habenula showed very low correlation (*r* < 0.10), suggesting a lack of functional connectivity between the bilateral habenula (Hétu et al., 2016). Distinct functional connectivity patterns of the left and right habenula with other subcortical and cortical regions have been repeatedly reported with data of variable quality (Erpelding et al., 2014; Hétu et al., 2016; Torrisi et al., 2017). Taken together, these findings support the functional laterality of the habenula in the human brain.

Regarding structural asymmetry of the habenula, only a few studies have been published, and these studies usually showed mixed results. For example, Ahumada-Galleguillos et al. (2017) found greater volume of the left habenula, in line with findings of multiple studies (e.g., Torrisi et al., 2017; Savitz et al., 2011b). However, other studies had inconsistent results: some studies did not find any inter-hemispheric differences in habenula volume (e.g., Lawson et al., 2013; Hétu et al., 2016), while others even suggested reversed asymmetry patterns (i.e., rightward) (e.g., Germann et al., 2020). These mixed results on habenular asymmetry may reflect differences in many factors, including sample heterogeneity such as age and sex distributions and sample sizes, as well as differences in brain data types (e.g., *in vivo* brain imaging vs. brain tissues), magnetic field strengths of the scanners (e.g., 3 T vs. 7 T), segmentation approaches (e.g., manual vs. automated), as we have observed in our large-scale cortical and subcortical asymmetry studies (Guadalupe et al., 2017; Kong et al., 2018; Kong et al., 2022; Ocklenburg & Güntürkün, 2018). Thus, a large-scale survey using meta-analysis approaches would be beneficial, to provide a clearer picture of the habenular asymmetry.

Moreover, altered asymmetry of the habenula has been suggested in various disorders. For example, a study using postmortem brain tissues from two groups of subjects with major and bipolar depression indicates the presence of asymmetric changes (a right-sided decrease) in habenular structure (Ranft et al., 2010). However, in some more recent studies, no significant differences were found in the volume of either the left or the right habenula (Luan et al., 2019; Schmidt et al., 2016). The left habenula showed a tendency towards lower volume in patients with schizophrenia (Zhang et al., 2017), while such differences were not found in another study (Ranft et al., 2010). As the results are mixed, a meta-analysis of existing data would be a major step forward in achieving a more accurate description of the variations related to these disorders.

In this study, we present a large-scale analysis of structural asymmetry of the habenula in the human brain, with the bilateral volume data from 52 datasets comprising 1,427 subjects, using meta-analytic methods. Our aim was to examine whether - and to what extent - the habenula shows left-right differences in volume, and to provide informative data on the habenular asymmetry with the largest sample size to date. We also estimated inter-dataset variability in the asymmetry effects, and assessed potential influences of age and sex on the variability in the asymmetry, as well as methodological factors such as MRI scanner field strength and the segmentation approaches used. Finally, we compared data on bilateral habenular volume and the left-right differences between patient datasets and controls, to test potential disorder-related effects.

## Materials and Methods

### Data sources

We searched PubMed, Web of Science, and Google Scholar for the articles that reported volume data on the bilateral habenula in the human brain, following PRISMA guidelines (Page et al., 2021). Specifically, the literature search was conducted on 19 April 2022, using the key terms: (“habenula volume*” OR “habenular volume*”) AND (human) AND (“magnetic resonance imaging” OR “MRI” OR “tissue”). Records covered the years from 1986 to 2022.

### Study selection

To screen the articles that might be relevant for the present study, we first removed duplicate records using EndNote, and two authors (Y.A. and X.K.) identified irrelevant entries by screening the titles and abstracts, and further retrieved the full-text of the remaining articles for data extraction (see below). Articles that did not report bilateral habenula volume data were excluded in the following analyses. Reference lists were also screened for additional references.

### Data extraction

From the selected articles, we extracted data on the mean and standard deviation (SD) of the left and right habenula volumes, various information on the samples (i.e., the sample sizes, mean ages and sex distributions of the samples [sex proportion of the sample, %male], and the sample category [e.g., healthy controls and major depressive disorder patients], and the measurement and segmentation approaches used). Note that healthy controls might include samples with other diseases that are not related to cerebral illness or neuropathy. For the majority of the articles, the data were accessible from the article text. In a few cases where the volume data was only presented within figures, we contacted the corresponding authors to request the exact numbers. When possible (Lawson et al., 2017; Liu et al., 2017; Germann et al., 2020), we extracted the volume data from these figures using an automatic tool, WebPlotDigitzer. Data from each study were double-extracted to prevent transcription errors. In addition, a study by Ranft et al. (2010) only reported the volume of the habenula subregions (i.e., the lateral and medial habenula). In this case, we combined the subregional data as an estimate of the whole habenula volume. When data were not accessible via any of these approaches, the studies were excluded from the follow-up statistical analyses.

During data extraction, we only kept one version of the data when datasets were identified as from the same or overlapping samples and were assessed using the same measurement approaches. Data from the overlapping samples but assessed using different measurement approaches were kept in the analyses for examining the effects of measurement approaches. For instance, we identified that the healthy individuals and the measurement approach used were identical in Savitz (2011a) and Savitz (2011b), so we only used that data once in the meta-analyses.

### Meta-analyses

We used mean difference (MD) and pooled standard deviation (SD_*puuled*_) derived from a dataset to assess the inter-hemispheric difference between the left and right habenula. Specifically, first the MD and pooled SD were calculated by following formulas (1) and (2) below, respectively. In the formula for calculating the pooled SD of each study, the correlation coefficient between the bilateral habenula volumes was taken into account for a closer estimation of the pooled SD. While such a correlation was not reported in the majority of these studies, we repeatedly used a range of various correlation values from 0.30 to 0.80 with a step size of 0.10 as suggested in recently study with a relatively large sample size (*N* > 100) (Germann et al., 2020). These derived measures were fed into a meta-analysis for estimating the left-right differences.

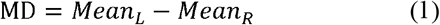

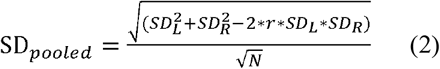

While the main analyses were based on data from all datasets, we repeated the analysis for the datasets from healthy controls, to reduce the heterogeneity that might be related to brain disorders. We also ran the same meta-analyses for the datasets from patients with one of the specific disorders included (i.e., major depressive disorders (MDD), schizophrenia (SCZ) and bipolar disorder (BD)).

Heterogeneity was tested using Cochran’s Q statistic, and quantified by the *I*^2^ statistic and *τ*^2^ statistic, which was calculated by the restricted maximum likelihood estimator (Viechtbauer, 2005). We also used Knapp-Hartung adjustments (Knapp & Hartung, 2003) to calculate the confidence interval around the pooled effect, to reduce the chance of false positives.

Given the similar results even with different correlations, in the main test, we focused on the results with a correlation of 0.50 (also discuss the results with other correlation values in the sensitivity analyses).

### Moderator analyses with meta-regression and subgroup analysis

To further address the source of heterogeneity in the meta-analyses, we investigated the potential effects of several moderating variables regarding the samples of each dataset, i.e., the sex ratio (%male) and mean age of the samples. In one case (Ahumada-Galleguillos et al., 2017), sex and age information were only available for the overall samples but not the subgroups, and thus the combined effect statistics were calculated. Other moderating factors were also considered, including brain data type (i.e., MRI images of post-mortem brain tissues), magnetic field strengths of the MRI scanners (i.e., 1.5 T, 3 T, and 7 T), and the segmentation approaches used (i.e., manual, fully automated, or semi-automated). In addition, we investigated a potential moderating effect of the average volume of bilateral habenula on the left-right differences. Meta-regression mixed effects models and subgroup analyses were used for the moderator analyses of continuous and categorical variables, respectively. When information for these variables were not available in some cases, the datasets were excluded from the corresponding analyses.

### Sensitivity analyses

The estimation of the pooled SD largely depends on the correlation coefficient of the left and right habenula volume, which in turn could impact the meta-analysis results. Thus, we repeated the analyses with a range of correlations (from 0.3 to 0.8), and reported the results in each case. In addition, including results based on too few participants may reduce reliability, so we re-ran the main analyses on datasets with a sample size of larger than 15.

### Software used

We used JASP (version 0.16) and R (version 4.2.0) for basic statistical analyses, and R package *meta* (version 5.2-0) for the meta-analyses, subgroup analysis, and meta-regressions.

## Results

### Study inclusion and data extraction

In total, we identified 244 records: 8, 12, and 222 from PubMed, Web of Science, and Google Scholar, respectively, and 2 additional records from reference lists (Fig. 2). After excluding duplicate records, 205 articles were left. Two authors (Y.A. and X.K.) read the abstract or full text as required, and removed 183 articles that did not report bilateral volume data for the habenula. One additional article was removed because of the duplicate sample issue that we identified (see Methods). Thus, 21 articles remained and were included in the main meta-analysis (Table 1).

**Table 1.**
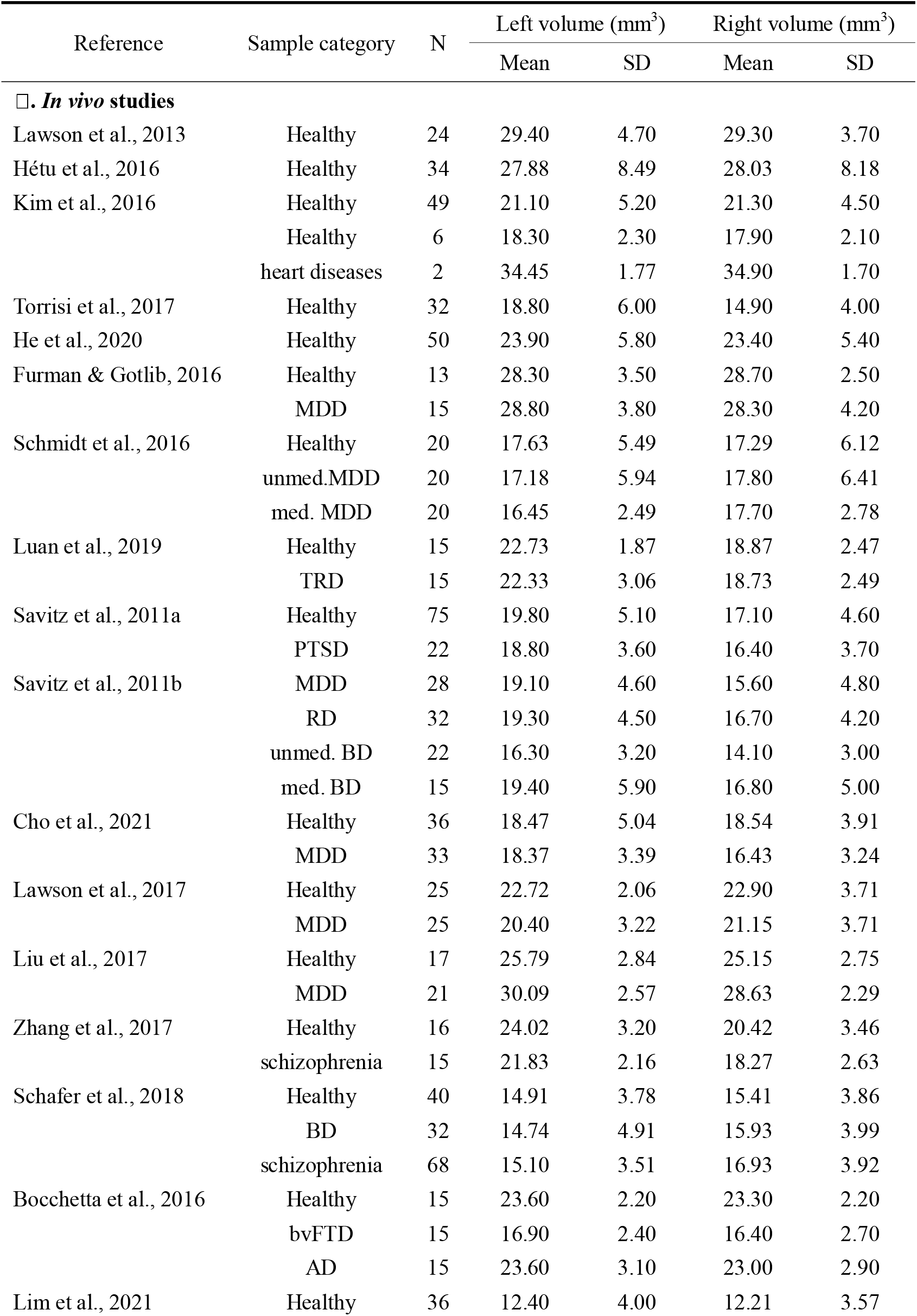
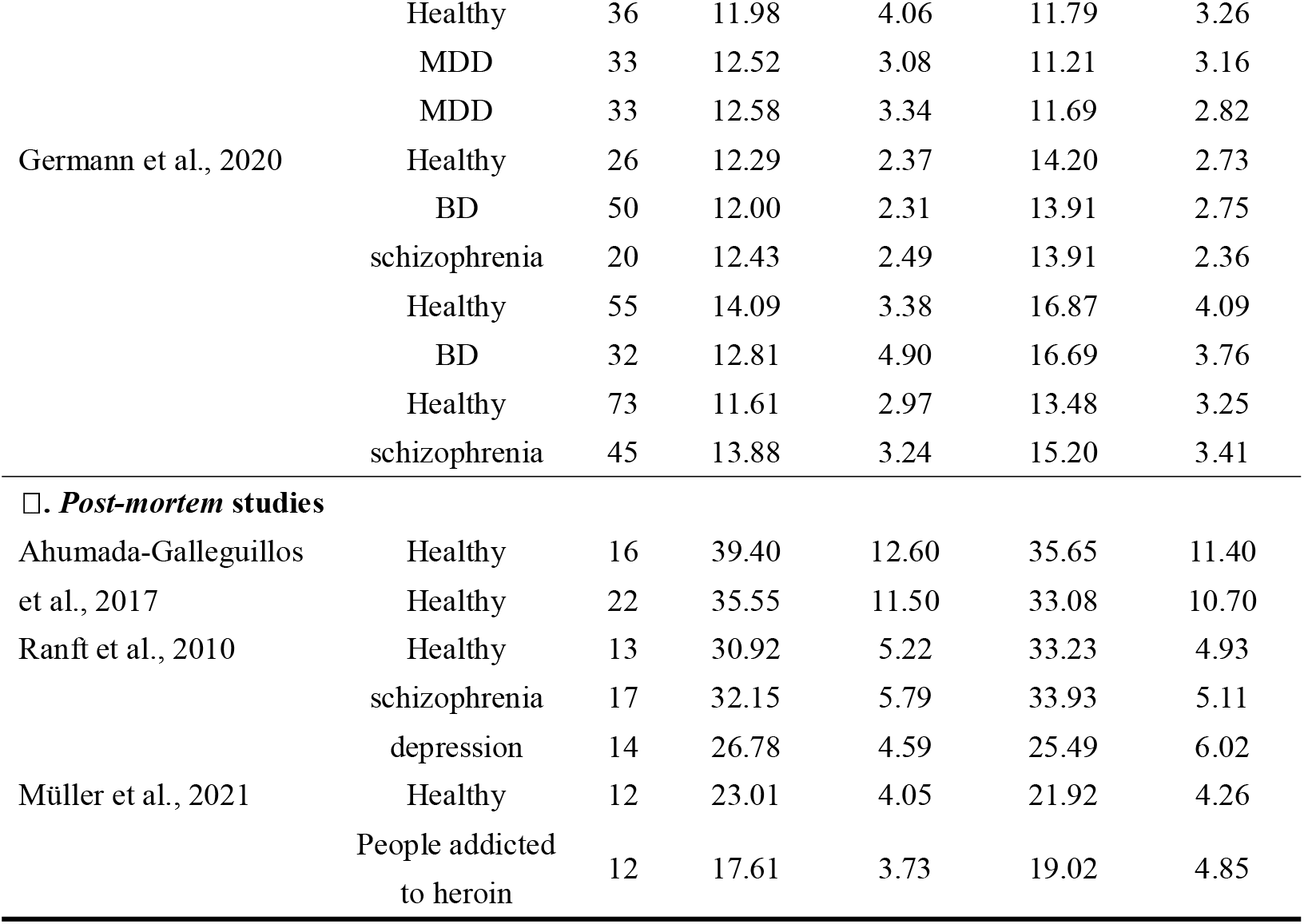
Data extracted from the habenula studies. *unmed*, unmedicated; *med*, medicated; MDD, major depressive disorder; TRD, treatment-resistant depression; PTSD, Post-traumatic stress disorder; RD, fully remitted patients with MDD; BD, bipolar disorder; bvFTD, behavioural variant frontotemporal dementia; AD, Alzheimer’s disease.

**Fig. 2:**
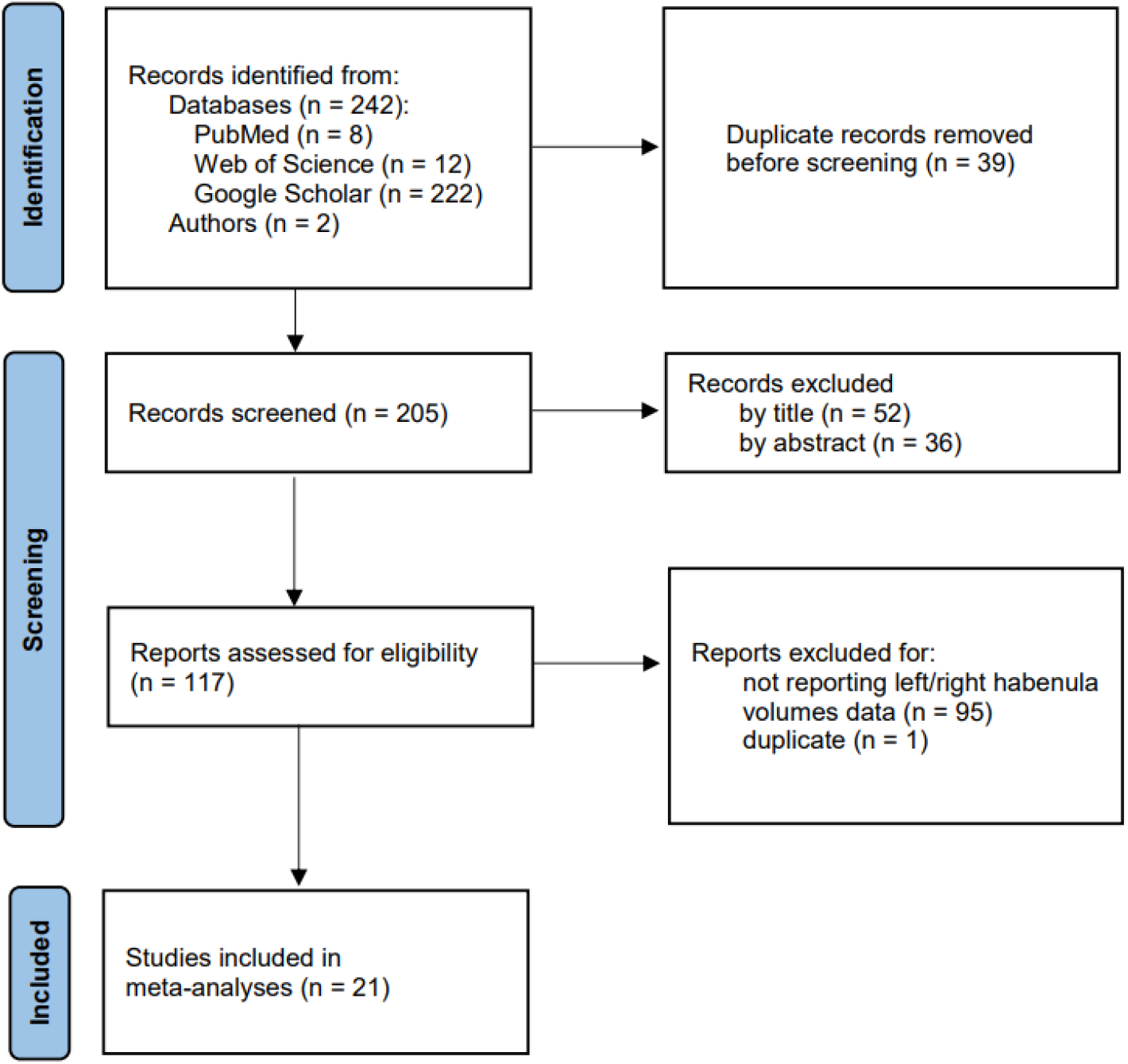
Flowchart of study selection.

Data on sample categories, sample size, and the mean and standard deviation of the bilateral habenula volume were extracted from each article (see Methods). This resulted in 52 separate datasets. These samples covered healthy individuals as well as patients with various diseases and conditions, including MDD, BD and SCZ. The sample sizes of the extracted volume data were up to 75 (mean = 27; median = 22; in total N = 1,427). Measurement approaches included brain imaging with 3 or 7 T MRI scanners, and image segmentation of brain tissues.

### Relationship between the volumes of the left and right habenula

The mean volume varied, ranging from 11.61 mm^3^ to 39.40 mm^3^, and from 11.21 mm^3^ to 35.65 mm^3^ for the left and right habenula, respectively, suggesting considerable variability of the habenula volume reported in the literature (Fig. 3). As expected, the left and right mean volume at the study level showed a high correlation when correlation was assessed across all the studies (*r* = 0.960, *p* < 0.001, number of datasets [k] = 52), as well as the various subgroups of healthy samples (*r* = 0.967, *p* < 0.001, k = 25), and patients with MDD (*r* = 0.957, *p* < 0.001, k = 11), SCZ (*r* = 0.960, *p* = 0.010, k = 5), but the correlation was relatively low in the BD patients (*r* = 0.361, *p* = 0.550, k = 5). These results suggested that little, if any, inter-hemispheric differences would exist in the habenula volume.

**Fig. 3:**
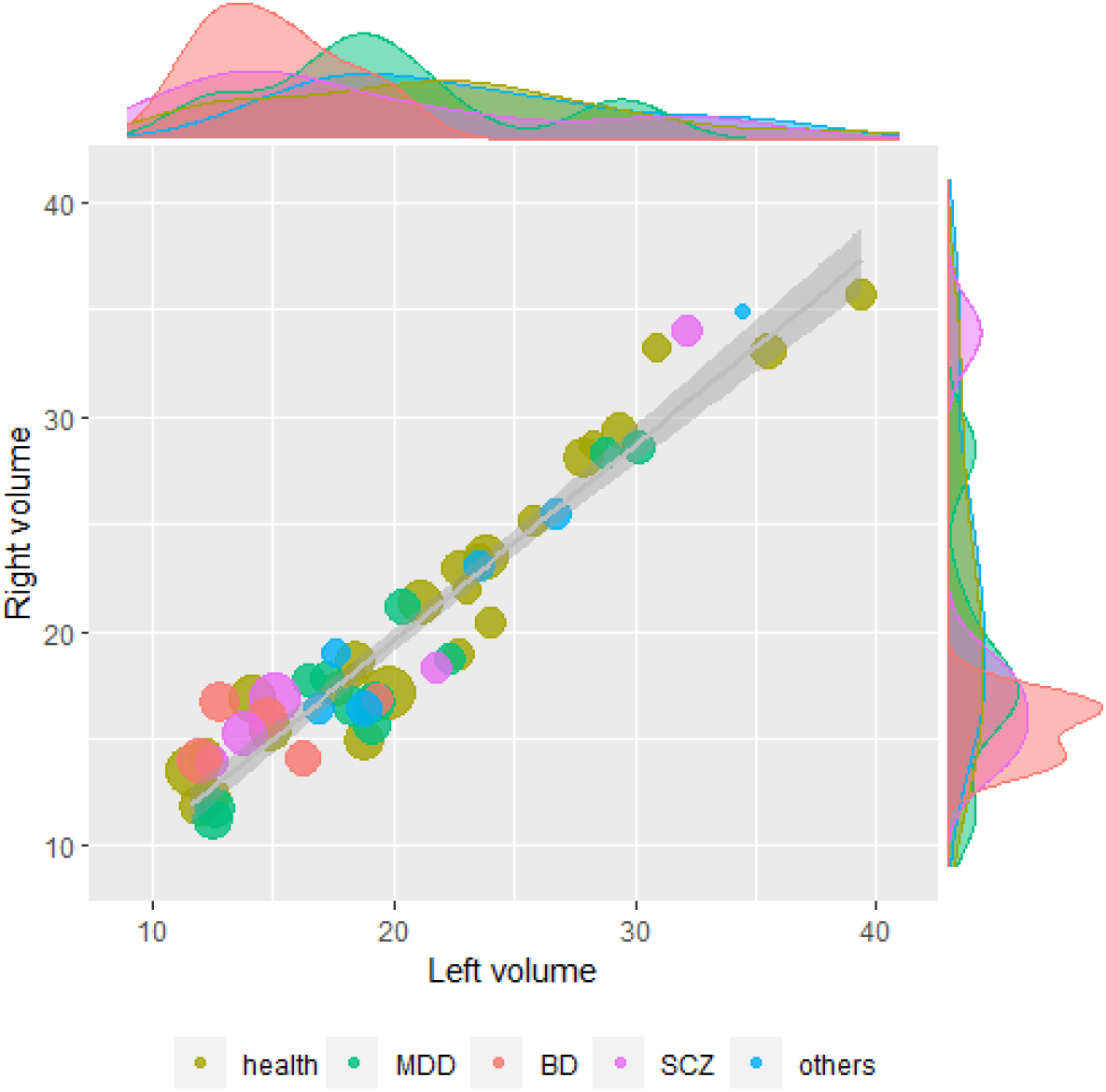
Scatter plot of the volume of the left (x-axis) and right (y-axis) habenula (mm^3^). Each dot indicates data from one dataset, with the size indicating its sample size and the color indicating the subgroups shown in the figure legend.

An initial synthesis of the volume data from the left and right habenula using a paired samples t-test also supported this point: no significant interhemispheric differences for either the healthy samples (*t*_(24)_ = 1.443, *p* = 0.162), or the patients with SCZ (*t*_(4)_ = -0.550, *p* = 0.612); there appeared to be a nominally significant difference in MDD subgroup (*t*_(10)_ = 2.402, *p* = 0.037), but it did not survive correction of multiple testing. In the BD subgroup, no significant differences (*t*_(4)_ = −0.352, *p* = 0.743) were observed, perhaps due to the limited number of studies. Note that this initial synthesis analysis was conducted at the study level, and potential individual-level variance and important variations across studies (e.g., sample size) were not considered. To further confirm these findings, a more formal synthesis was conducted using a meta-analysis approach below.

### Inter-hemispheric differences in the habenula volume

#### Meta-analysis of all datasets

To take into account individual-level variances and the various samples of these studies, we performed a meta-analysis by integrating the MD and pooled SD data. The habenula showed positive MDs in 30 datasets (maximum MD: 3.90 mm^3^_;_ Left > Right; *N* = 720), and negative MDs in the remaining 22 datasets (MD of the maximum difference: -3.88 mm^3^; Left < Right; *N* = 707). The meta-analysis of the left-right differences in all the datasets showed an average difference of 0.41 mm^3^, with a 95% confidence interval (CI) of −0.13 mm^3^ to 0.94 mm^3^. This observed difference was not significant with a significance threshold of p < 0.05 (t_(51)_= 1.52, p = 0.134) (Fig.4).

**Fig. 4:**
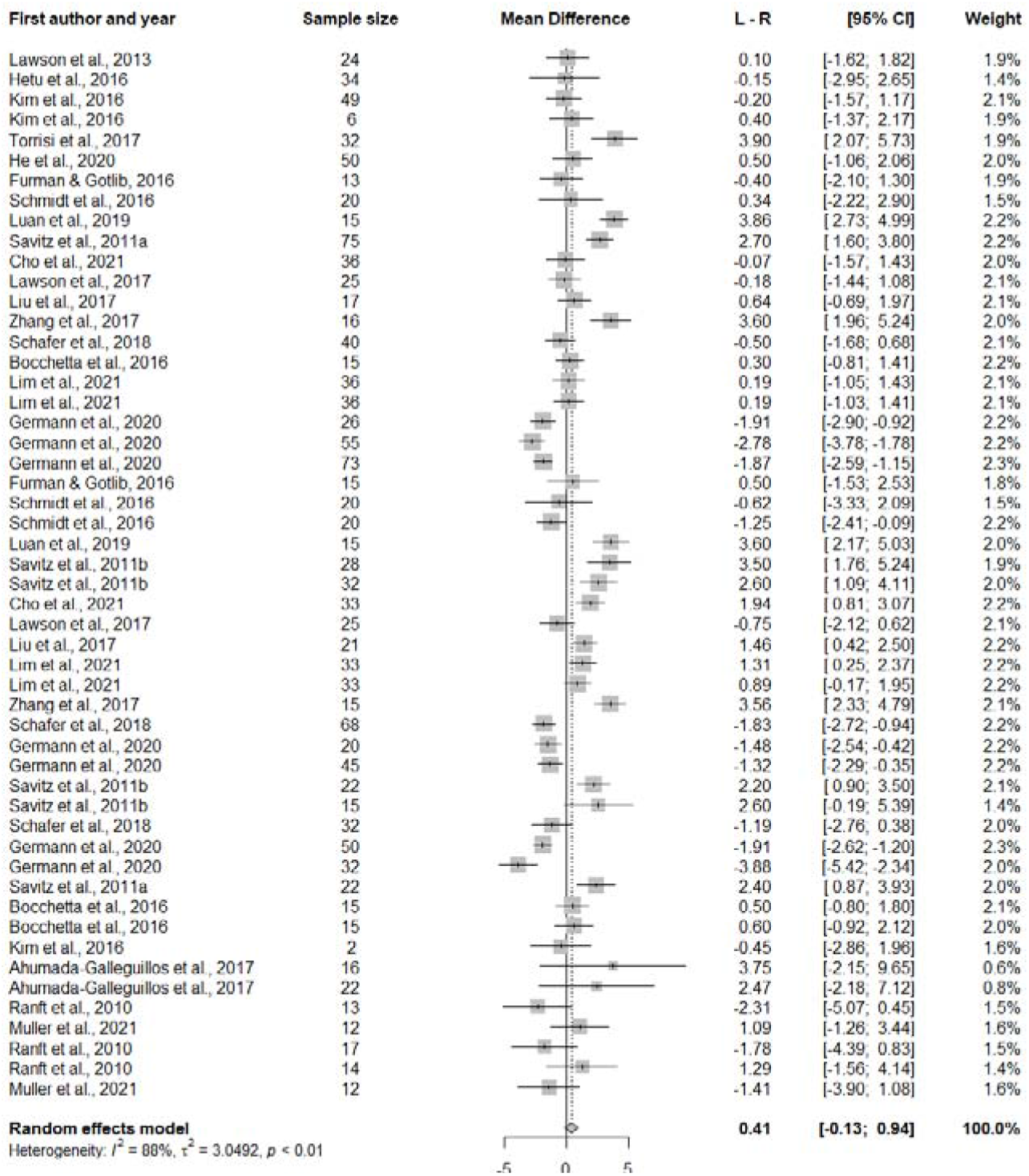
Forest plot of the left-right differences in the habenula volume.

#### Meta-analysis of different groups of patients

We also ran meta-analyses of the inter-hemispheric differences in the volumes for the datasets of healthy samples and each patient group (i.e., patients with MDD, SCZ and BD; Fig. 5). In line with the overall findings above, no differences were found in either of these groups: healthy (MD = 0.40 mm^3^; 95% CI: −0.37 mm^3^ to 1.16 mm^3^; *t*_(24)_ = 1.07, *p* = 0.294), SCZ (MD = −0.54 mm^3^_;_ 95% CI: -3.44 mm^3^ to 2.36 mm^3^; *t*_(4)_ < 1.00), BD (MD = −0.54 mm^3^; 95% CI: -3.94 mm^3^ to 2.85 mm^3^; *t*_(4)_ < 1.00). Interestingly, the meta-analysis also showed a significant left-right difference in the MDD samples (MD = 1.23 mm^3^; 95% CI: 0.14 mm^3^ to 2.33 mm^3^; Left > Right: *t*_(10)_ = 2.51, *p* = 0.031), which is similar to that of the initial comparison above. We did not find any significant differences in the left-right difference of the habenula volume between either of the disorder groups and the healthy control samples (*p*s > 0.05).

**Fig. 5:**
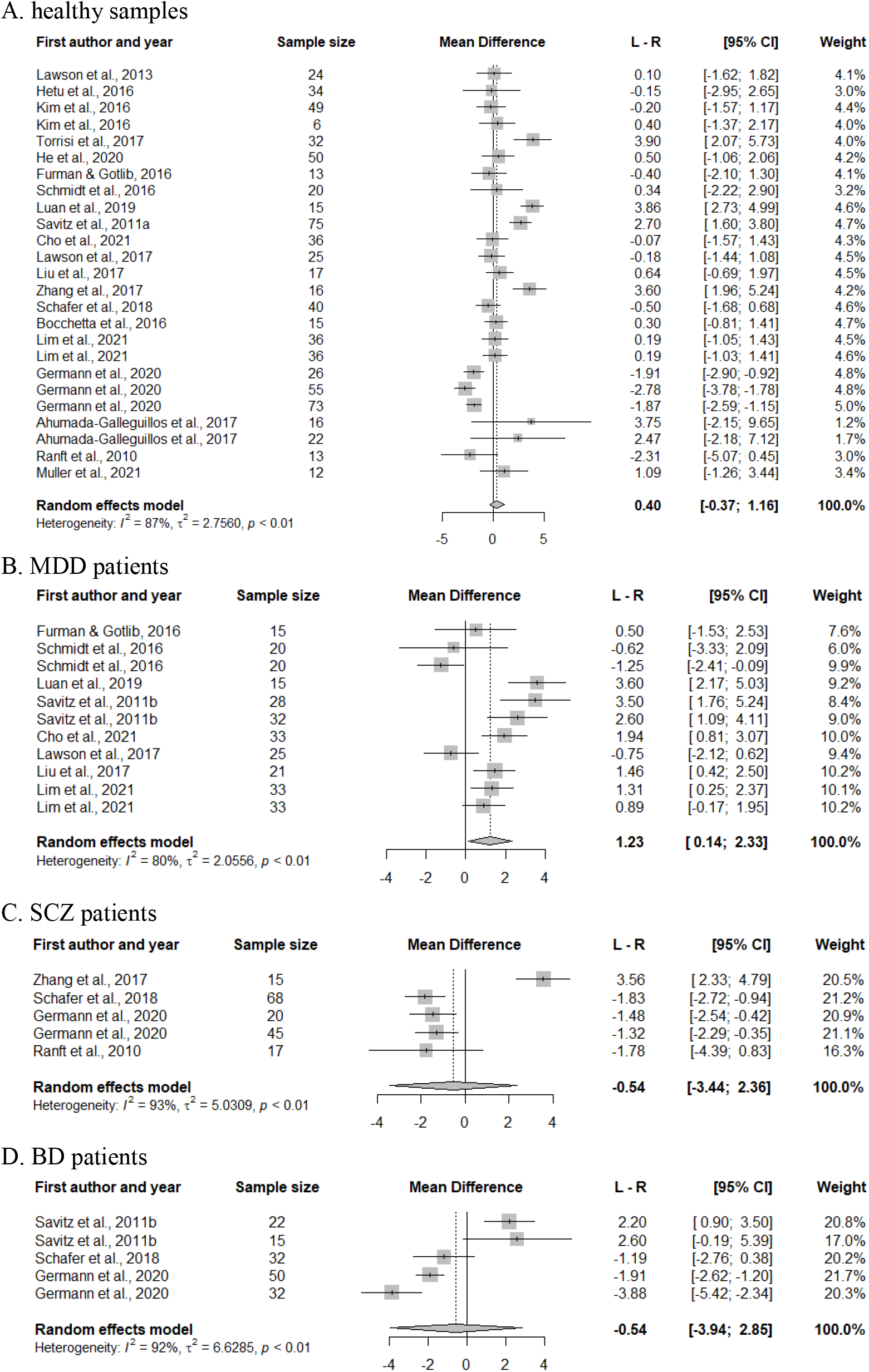
Forest plots of inter-hemispheric habenula volume differences in different subgroups.

### Moderator analyses

Moderate to substantial heterogeneity in the MD was observed across the datasets. The heterogeneity remained when analyzing only datasets from either the healthy controls (*I*^2^ = 87%, τ^2^ =2.756, *p* < 0.01), or specific disorder samples (MDD: *I*^2^ = 80%, τ^2^ = 2.056, *p* < 0.01; SCZ: *I*^2^ = 93%, τ^2^ = 5.031, *p* < 0.01; BD: *I*^2^ = 92%, τ^2^ = 6.629, *p* < 0.01). To further address the heterogeneity in the meta-analyses, we investigated the potential effects of several moderating variables, including the sex ratio and age of samples in each dataset, brain data type (*in vivo* versus *post mortem*), magnetic field strengths of the MRI scanners, and the segmentation approaches used.

#### Effects of sex ratio and mean age

Information on the sex distribution and mean age of the samples was available for 50 and 42 datasets, respectively. Meta-regression analyses with either sex ratio or mean age as moderators showed no significant effects (Sex: *R*^2^ = 5.25%, *F*_(1, 40)_ = 2.81, *p* = 0.102; Age: *R*^2^ = 5.54%, *F*_(1, 48)_ = 2.81, *p* = 0.100). We also repeated the analyses within the healthy control datasets, and did not find any significant effects (Sex: *p* = 0.961; Age: *p* = 0.570) (given the limited number of datasets available, we did not run separate moderator analyses for each patient group, the same as below).

#### Effects of brain data type

In terms of brain data types used for the measurement of the habenula volume, 44 datasets were based on *in vivo* studies (i.e., using MRI), and 8 were based on *post-mortem* studies (i.e., via brain tissues). Subgroup analysis showed no significant differences associated with the brain data types (χ^2^_(1)_ = 1.07, *p* = 0.302), and the analysis with only the healthy control datasets showed similar results (χ^2^ = 0.04, *p* = 0.850).

#### Effects of the magnetic field strength of the MRI scanners

Within the samples using MRI, we further investigated the effects of various MRI scanners. Here we focused on the magnetic field strength of the scanners, which is perhaps the main factor that could affects the brain imaging measures. Among the 44 datasets, 7 were based on 1.5 T MRI scanners, 26 were based on 3 T, and 11 were based on 7 T (Fig. 6). As expected, subgroup analyses showed significant moderating effects (χ^2^_(2)_ = 65.37, p < 0.001), - effects that were mainly contributed by the significant left-right differences observed in the 3 T datasets (MD = 1.15 mm^3^; 95%CI: 0.45 to 1.84; left > right: *t*_(25)_ = 3.41, *p* = 0.002) and 1.5 T datasets (MD = -2.02 mm^3^; 95%CI: -2.66 to −1.37; left < right: *t*_(6)_ = -7.63, *p* = 0.0003). With only the healthy control datasets, the results were similar in the subgroup analyses of both 3 T scanners (MD = 0.90mm^3^; 95%CI: −0.12 to 1.92; left > right: *t*_(11)_ =1.95, *p* = 0.077; and 1.5 T scanners (MD = -2.12mm^3^; 95%CI: -3.31 to −0.92; left < right: *t*_(2)_ = -7.62, *p* = 0.017). Note that the samples of the 3 T and 1.5 T datasets showed inversed asymmetry patterns which might also relate to different measurement approaches used such as segmentation (see below and also Discussion). In addition, no significant differences were found in the 7 T datasets (*t*_(10)_ = 1.68, *p* = 0.125).

**Fig. 6:**
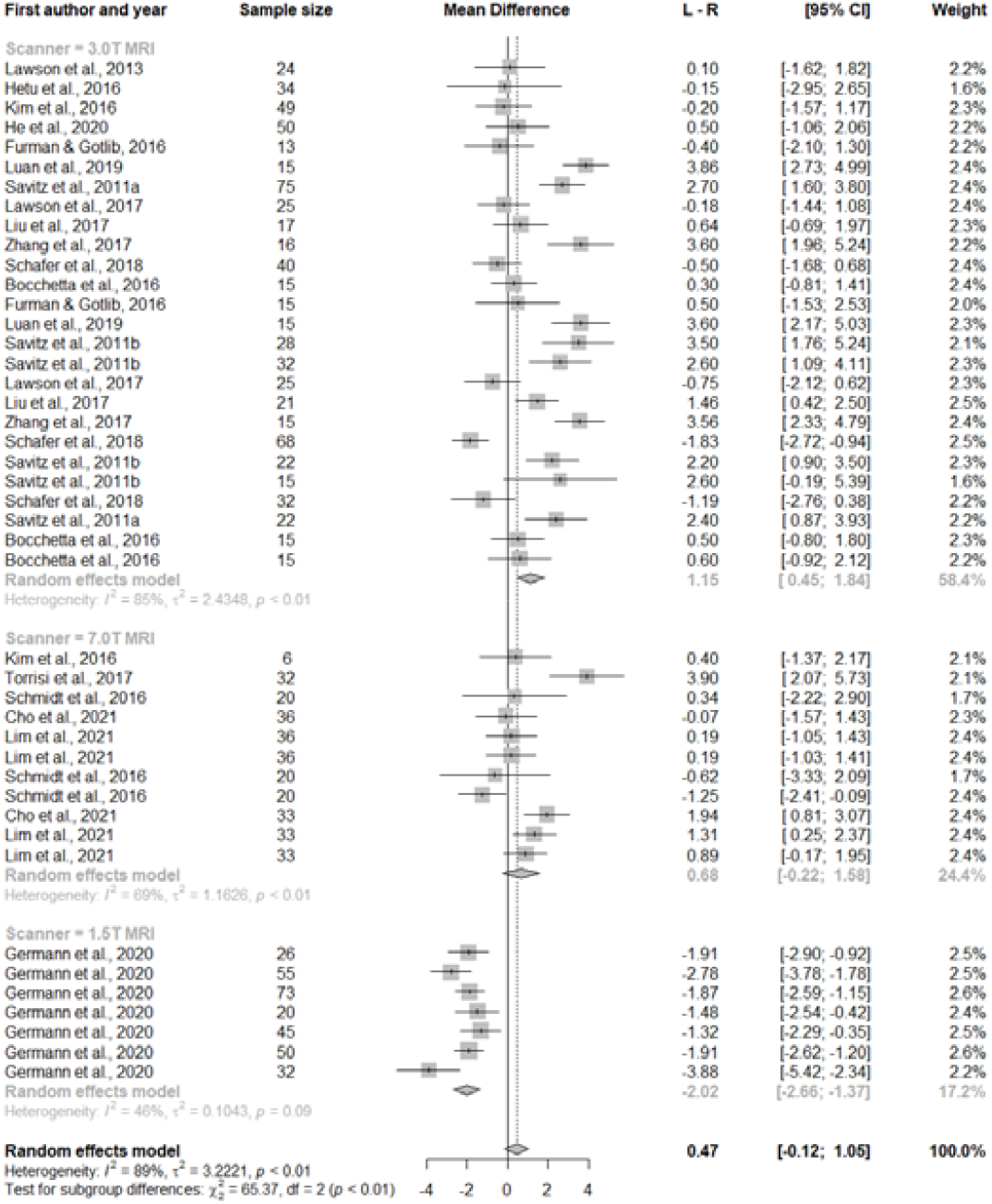
Forest plots for the moderator analysis of magnetic field strength of the MRI scanners.

#### Effects of segmentation methods

The various segmentation methods used could be another vital factor contributing to the observed heterogeneity. Among the 44 MRI datasets, 28 were identified as using manual segmentation approaches, 9 used fully automated segmentation, and 7 used semi-automated segmentation (Fig. 7). Subgroup analyses by segmentation method showed significant moderating effects (χ^2^_(2)_ = 52.32, p < 0.001). Specifically, we found that in the manual subgroup, the habenula volume was significantly larger in the left hemisphere compared to the right (MD = 1.49 mm^3^; 95% CI: 0.91 to 2.07; left > right: *t*_(27)_ = 5.27, *p* < 0.0001), while inverse patterns were found in the semi-automated (MD = −0.99; 95% CI: −1.65 to −0.33; left < right: *t*_(6)_ = -3.68, *p* = 0.010) and fully-automated subgroups (MD = −1.54; 95% CI: -2.61 to −0.48; left < right: *t*_(8)_ = -3.35, *p* = 0.010).

**Fig. 7:**
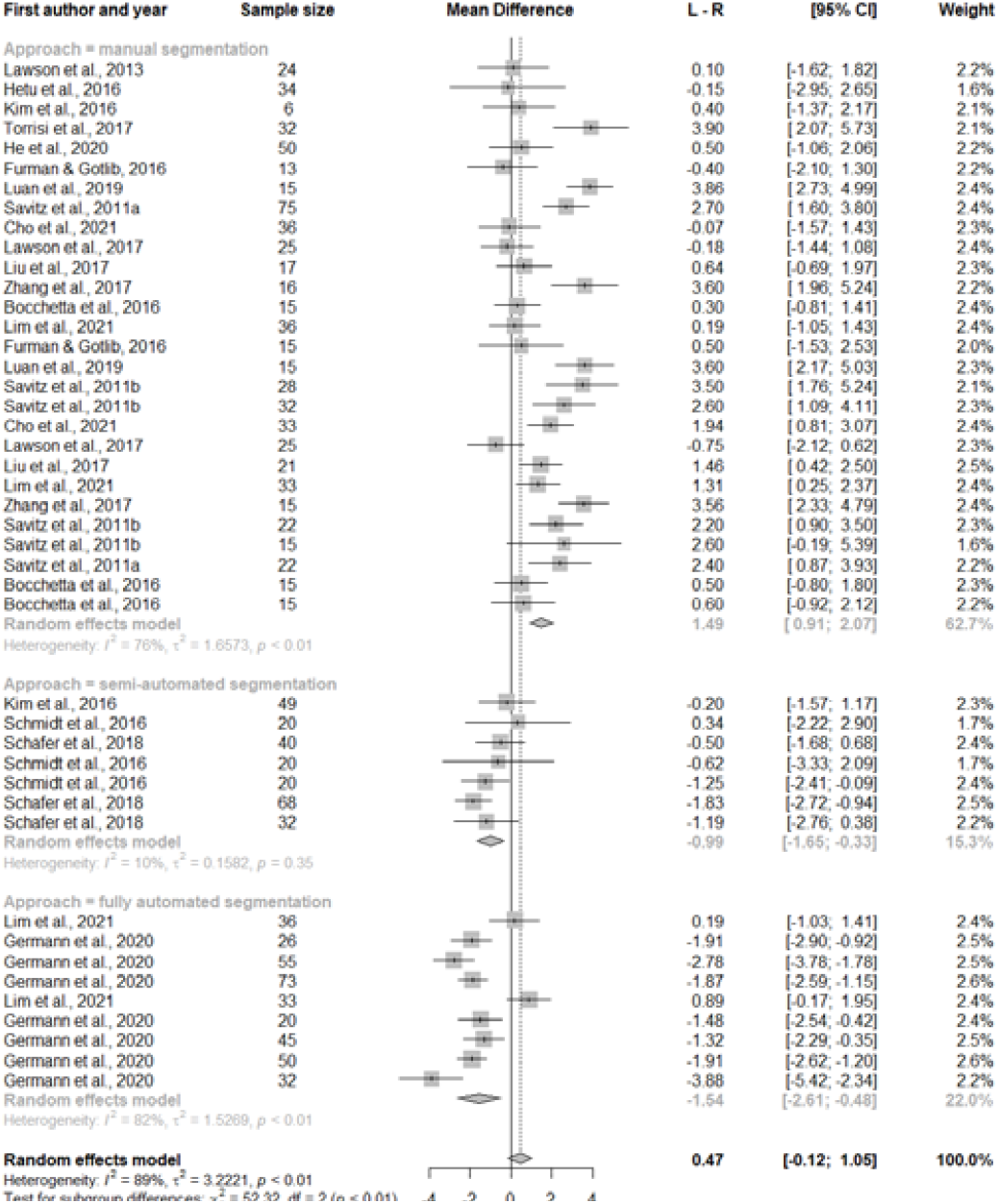
Forest plots for the moderator analysis of segmentation approaches.

#### Effects of the average volume of the bilateral habenula

We also investigated the moderating effects of the volume of the habenula on the left-right difference estimation. Meta-regression analyses showed no significant effects for either analysis across all the datasets, or for that within the healthy control datasets (all: *R*^2^ = 1.10%, *F*(1, 50) = 1.11, *p* = 0.298; healthy: *R*^2^ = 0.45%, *F*(1, 23) = 0.96, *p* = 0.338).

### Sensitivity analyses

To rule out the potential confounding effects of the correlation of the bilateral habenula volume on the meta-analysis results, we repeated all analyses with a range of correlation values from 0.30 to 0.80. The results remained similar in all cases (Fig. 8).

**Fig. 8:**
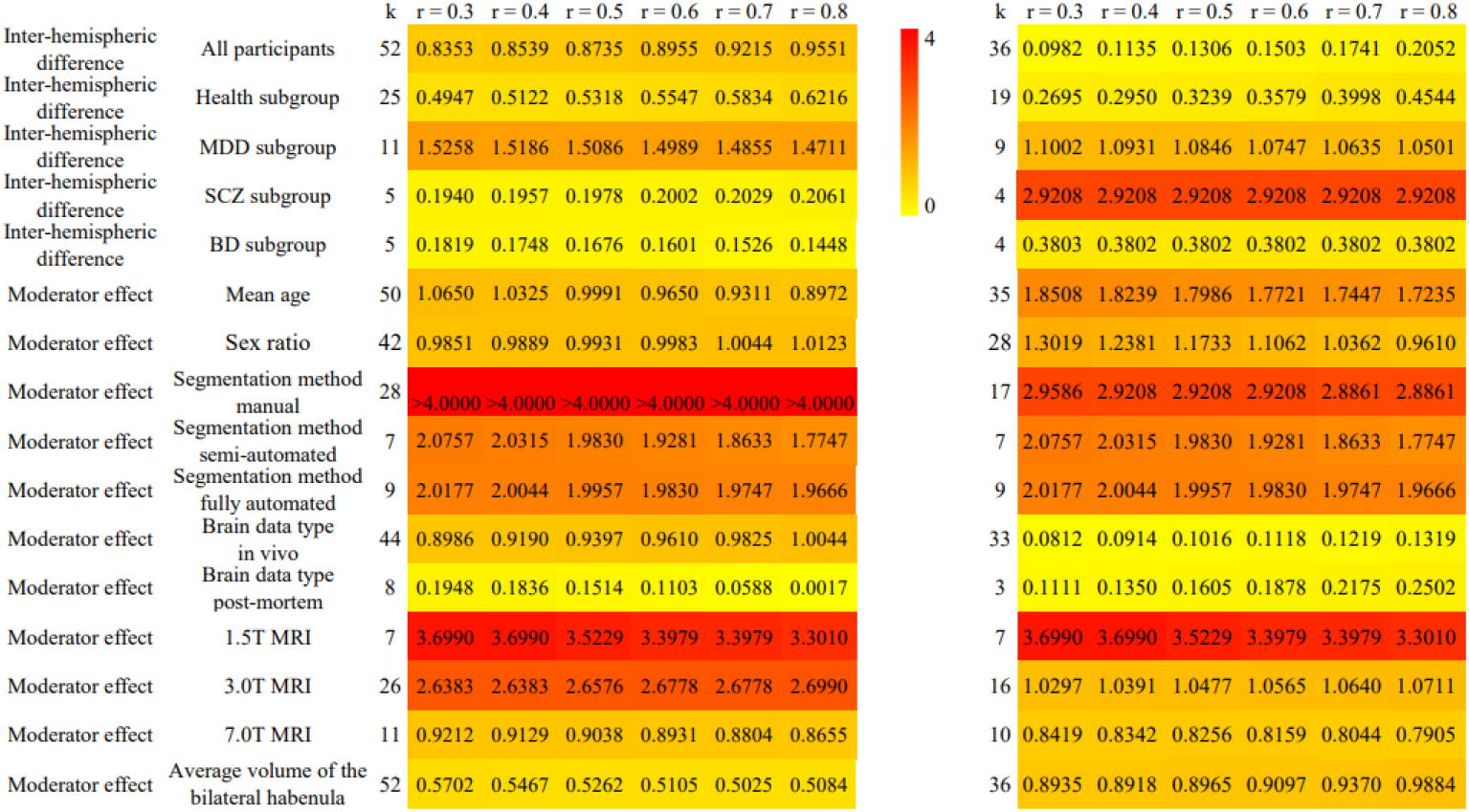
Results of the sensitivity analyses. Color indicates the statistical significance (*-log(p)*) in each analysis: the left panel for analyses with all dataset, the right for analyses after excluding the datasets with no more than 15 samples.

In addition, to evaluate the potential effects of the datasets with too few samples in the meta-analyses, we excluded these small datasets (i.e., those with no more than 15 individuals) and re-ran all analyses. The number of datasets included in the meta-analyses decreased remarkably, at least in some cases, but the main results remained similar except for the subgroup analyses of the left-right differences in the SCZ samples (Fig. 8). Specifically, the inter-hemispheric differences in the SCZ samples became highly significant after excluding one small dataset (*p* < 0.002).

### Disorder-related differences in the habenula volume

We also used the data generated to investigate disorder-related differences in habenular volume. Meta-analyses showed no significant differences in the unilateral habenula volume of either samples with MDD (left: *t*_(10)_ = 0.01; right: *t*_(10)_ = −0.53), SCZ (left: *t*_(4)_ = 0.43; right: *t*_(4)_ = 0.57), or BD (left: *t*_(4)_ = -2.07, *p* = 0.107; right: *t*_(4)_ = −1.08, *p* = 0.341), compared to healthy control datasets from the same study. Cross-disorder comparisons showed no significant differences (*p*s > 0.05) except for the differences in the left habenula between the MDD subgroup and the BD subgroup (MDD > BD; *Q*_*(1)*_ = 7.70, *p* = 0.006).

## Discussion

This study presents a large-scale analysis of structural asymmetry of the habenula in the human brain. Meta-analyses of 52 datasets (*N* = 1,427) revealed significant heterogeneity in the left-right differences in the habenula volume across datasets. The magnetic field strength of the MRI scanners (e.g., 3 T or 1.5 T) and segmentation methods (e.g., manual or automated) applied were found to be two of the key contributing factors. While little evidence was found for volume asymmetry in the meta-analysis of all of the datasets, the most pronounced left-right differences (i.e., left > right) were found in more homogeneous subgroup of datasets with 3 T scanners and the manual segmentation approach. We did not find significant disorder-related differences, relative to healthy controls in either the left-right differences or the unilateral volume *per se*.

### Left-right differences in the volume of the habenula

The habenula volume of the left and right hemispheres showed a high correlation (*r* = 0.96) across datasets (i.e., at the study level), suggesting high cross-hemisphere similarity for the bilateral structure in the human brain. This seems to be inconsistent with the correlations reported at the individual level (around 0.50) (e.g., Germann et al., 2020), which suggests a considerable extent of distinctness. Such inconsistency could be due to higher measurement errors of individual data, and/or larger variance across studies at the study-level analysis. Moreover, mixed asymmetry patterns of the habenula volume have been suggested in the previous studies (see Introduction). The present study, with a large number of samples, revealed little evidence to support left-right differences in the habenula volume. Interestingly, we successfully identified several significant moderating factors that could impact the left-right comparison results, such as the field strength of the scanners and the segmentation approaches. In terms of scanners, we found that datasets from 3 T scanners tended to show a leftward asymmetry in the habenular volume, while datasets from 1.5 T scanners tended to show an inverse pattern (the 1.5 T datasets were from the same study, i.e., Germann et al., 2020). Datasets from 7 T scanners showed no significant asymmetry although we would expect better imaging quality with such a high field strength. The scanner effects seem to be unique to the habenular volume asymmetry as we did not find significant effects on the asymmetry of cortical thickness or area (Kong et al., 2018), or the volume asymmetry of larger subcortical structures such as the hippocampus or thalamus (Guadalupe et al., 2017). In terms of segmentation approaches, MRI studies using a manual approach tended to report a leftward asymmetry, while those using a semi- or fully-automated approach tended to report a rightward asymmetry. There could be some interesting interactions between scanner-related factors and the segmentation approaches used in measuring the hemisphere-related differences, which could be addressed in future studies.

To our knowledge, this is the first large-scale empirical investigation of habenular structural asymmetry. The results provided robust data on the moderating effects of multiple factors including scanners and habenula segmentation. Such moderating effects suggested that when developing new segmentation algorithms for future studies, the potential asymmetrical nature of the brain structures should be taken into consideration if possible.

### Disorder-related differences in the habenula

The habenula has been implicated in a variety of psychiatric disorders such as depression and anxiety disorders (Boulos et al., 2017; Fakhoury, 2017), and also considered as a promising target for the treatment of intractable conditions (Sartorius et al., 2007; 2010). A few brain structural studies have suggested asymmetrical alterations in the habenular volume in patients with e.g., MDD (e.g., Ranft et al., 2010) or SCZ (e.g., Zhang et al., 2017). However, the present meta-analyses showed little differences when comparing the unilateral habenular volume data of patients with that of the healthy controls. Interestingly, we found that patients with MDD tended to show a larger volume in the left hemisphere than the right, and patients with SCZ seemed to possess a larger habenular volume in the right hemisphere. Note that these findings were obtained using a relatively smaller number of datasets and meanwhile could be confounded by variability in brain imaging approaches (e.g., all datasets with 7T or 3T scanners for MDD versus half of the datasets with 1.5 T scanners for SCZ). Thus such differences should be interpreted with caution. At best, these observations could serve as potential hypotheses for additional experiments that include independent data.

### Limitations and future directions

This study has several limitations that could be overcome in future studies. First, while this is to our best knowledge the largest study on the habenular asymmetry, the findings warrant further investigation with a larger number of individuals. The UK Biobank cohort (Sudlow et al., 2015) and the ENIGMA working groups (Thompson et al., 2020) could provide great opportunities for achieving a more definitive picture of the nature of human habenula asymmetry. Second, an automated and unbiased segmentation approach is necessary for analyzing such large-scale individual data. Currently, a few approaches have been proposed such as MAGeTbrain (Pipitone et al., 2014; Chakravarty et al., 2013), myelin content-based segmentation (Kim et al., 2016; Kim et al., 2018), and deep learning-based U-Net segmentation (Lim et al., 2021). It remains to be determined whether such approaches could detect any hemispheric differences in the habenula. In addition, the habenula can be divided into medial and lateral habenula both structurally and functionally in animal studies (Fakhoury & Domínguez López, 2014; Hikosaka, 2010). Ultra-high-resolution *in vivo* MR imaging could provide new data on the functional lateralization of the habenula in the human brain and its structural basis.

### Summary

In summary, the present study presents a large-scale analysis of structural asymmetry of the habenula in the human brain. Results showed significant heterogeneity in both the habenula volume and its left-right differences across different studies. The magnetic field strength of the MRI scanners and segmentation methods used were found to be two of the key contributing factors to such heterogeneity. The most pronounced left-right differences (i.e., left > right) were found in datasets with 3 T scanners and manual segmentation approach. While inversed asymmetry patterns were suggested in MDD (left > right) and SCZ (left < right) patients, little evidence for the differences was found when comparing with healthy samples or other disorder samples. This study provides useful data for future studies of brain imaging and methodological developments related to precision habenula segmentation, and also contributes to the understanding of potential roles of habenular laterality in health and disorders.

## Acknowledgments

Xiang-Zhen Kong is supported by the Fundamental Research Funds for the Central Universities (2021XZZX006), the National Natural Science Foundation of China (32171031), and Information Technology Center of Zhejiang University. Paul M. Thompson is supported in part by the U.S. National Institutes of Health under grant R01 MH116147. We thank Prof. Clyde Francks (Language and Genetics Department, Max Planck Institute for Psycholinguistics, Netherlands) for helpful discussion and comments on the manuscript.

## Data and code availability

Our study primarily relies on the dataset extracted from the published papers. We provide the data and code at [link to be updated upon acceptance].

## Author contributions

Y.A.: study conception and design, data preparation, analysis, visualization, and preparing the first draft; Y.P. and P.M.T.: writing and editing; X.K.: study conception and design, data preparation, analysis, visualization, preparing the first draft and editing.

## References

Agetsuma, M., Aizawa, H., Aoki, T., Nakayama, R., Takahoko, M., Goto, M., Sassa, T., Amo, R., Shiraki, T., Kawakami, K., Hosoya, T., Higashijima, S., & Okamoto, H. (2010). The habenula is crucial for experience-dependent modification of fear responses in zebrafish. Nature Neuroscience, 13(11), 1354–1356. https://doi.org/10.1038/nn.2654

Ahumada-Galleguillos, P., Lemus, C. G., Díaz, E., Osorio-Reich, M., Härtel, S., & Concha, M. L. (2017). Directional asymmetry in the volume of the human habenula. Brain Structure and Function, 222(2), 1087–1092. https://doi.org/10.1007/s00429-016-1231-z

Amo, R., Aizawa, H., Takahoko, M., Kobayashi, M., Takahashi, R., Aoki, T., & Okamoto, H. (2010). Identification of the Zebrafish Ventral Habenula As a Homolog of the Mammalian Lateral Habenula. Journal of Neuroscience, 30(4), 1566–1574. https://doi.org/10.1523/JNEUROSCI.3690-09.2010

Batalla, A., Homberg, J. R., Lipina, T. V., Sescousse, G., Luijten, M., Ivanova, S. A., Schellekens, A. F. A., & Loonen, A. J. M. (2017). The role of the habenula in the transition from reward to misery in substance use and mood disorders. Neuroscience & Biobehavioral Reviews, 80, 276–285. https://doi.org/10.1016/j.neubiorev.2017.03.019

Bocchetta, M., Gordon, E., Marshall, C. R., Slattery, C. F., Cardoso, M. J., Cash, D. M., Espak, M., Modat, M., Ourselin, S., Frisoni, G. B., Schott, J. M., Warren, J. D., & Rohrer, J. D. (2016). The habenula: An under-recognised area of importance in frontotemporal dementia? Journal of Neurology, Neurosurgery & Psychiatry, 87(8), 910–912. https://doi.org/10.1136/jnnp-2015-312067

Boulos, L.-J., Darcq, E., & Kieffer, B. L. (2017). Translating the Habenula—From Rodents to Humans. Biological Psychiatry, 81(4), 296–305. https://doi.org/10.1016/j.biopsych.2016.06.003

Chakravarty, M. M., Steadman, P., van Eede, M. C., Calcott, R. D., Gu, V., Shaw, P., Raznahan, A., Collins, D. L., & Lerch, J. P. (2013). Performing label-fusion-based segmentation using multiple automatically generated templates: MAGeT Brain: Label Fusion Segmentation Using Automatically Generated Templates. Human Brain Mapping, 34(10), 2635–2654. https://doi.org/10.1002/hbm.22092

Cho, S.-E., Park, C.-A., Na, K.-S., Chung, C., Ma, H.-J., Kang, C.-K., & Kang, S.-G. (2021). Left-right asymmetric and smaller right habenula volume in major depressive disorder on high-resolution 7-T magnetic resonance imaging. PLOS ONE, 16(8), e0255459. https://doi.org/10.1371/journal.pone.0255459

Dorsaint-Pierre, R., Penhune, V. B., Watkins, K. E., Neelin, P., Lerch, J. P., Bouffard, M., & Zatorre, R. J. (2006). Asymmetries of the planum temporale and Heschl’s gyrus: Relationship to language lateralization. Brain, 129(5), 1164–1176. https://doi.org/10.1093/brain/awl055

Dreosti, E., Vendrell Llopis, N., Carl, M., Yaksi, E., & Wilson, S. W. (2014). Left-Right Asymmetry Is Required for the Habenulae to Respond to Both Visual and Olfactory Stimuli. Current Biology, 24(4), 440–445. https://doi.org/10.1016/j.cub.2014.01.016

Duboc, V., Dufourcq, P., Blader, P., & Roussigné, M. (2015). Asymmetry of the Brain: Development and Implications. Annual Review of Genetics, 49(1), 647–672. https://doi.org/10.1146/annurev-genet-112414-055322

Erpelding, N., Sava, S., Simons, L. E., Lebel, A., Serrano, P., Becerra, L., & Borsook, D. (2014). Habenula functional resting-state connectivity in pediatric CRPS. Journal of Neurophysiology, 111(2), 239–247. https://doi.org/10.1152/jn.00405.2013

Eyler, L. T., Pierce, K., & Courchesne, E. (2012). A failure of left temporal cortex to specialize for language is an early emerging and fundamental property of autism. Brain, 135(3), 949–960. https://doi.org/10.1093/brain/awr364

Fakhoury, M. (2017). The habenula in psychiatric disorders: More than three decades of translational investigation. Neuroscience & Biobehavioral Reviews, 83, 721–735. https://doi.org/10.1016/j.neubiorev.2017.02.010

Fakhoury, M., & Domínguez López, S. (2014). The Role of Habenula in Motivation and Reward. Advances in Neuroscience, 2014, 1–6. https://doi.org/10.1155/2014/862048

Furman, D. J., & Gotlib, I. H. (2016). Habenula responses to potential and actual loss in major depression: Preliminary evidence for lateralized dysfunction. Social Cognitive and Affective Neuroscience, 11(5), 843–851. https://doi.org/10.1093/scan/nsw019

Germann, J., Gouveia, F. V., Martinez, R. C. R., Zanetti, M. V., de Souza Duran, F. L., Chaim-Avancini, T. M., Serpa, M. H., Chakravarty, M. M., & Devenyi, G. A. (2020). Fully Automated Habenula Segmentation Provides Robust and Reliable Volume Estimation Across Large Magnetic Resonance Imaging Datasets, Suggesting Intriguing Developmental Trajectories in Psychiatric Disease. Biological Psychiatry: Cognitive Neuroscience and Neuroimaging, 5(9), 923–929. https://doi.org/10.1016/j.bpsc.2020.01.004

Guadalupe, T., Mathias, S. R., vanErp, T. G. M., Whelan, C. D., Zwiers, M. P., Abe, Y., Abramovic, L., Agartz, I., Andreassen, O. A., Arias-Vásquez, A., Aribisala, B. S., Armstrong, N. J., Arolt, V., Artiges, E., Ayesa-Arriola, R., Baboyan, V. G., Banaschewski, T., Barker, G., Bastin, M. E., … Francks, C. (2017). Human subcortical brain asymmetries in 15,847 people worldwide reveal effects of age and sex. Brain Imaging and Behavior, 11(5), 1497–1514. https://doi.org/10.1007/s11682-016-9629-z

Guadalupe, T., Willems, R. M., Zwiers, M. P., Arias Vasquez, A., Hoogman, M., Hagoort, P., Fernandez, G., Buitelaar, J., Franke, B., Fisher, S. E., & Francks, C. (2014). Differences in cerebral cortical anatomy of left- and right-handers. Frontiers in Psychology, 5. https://doi.org/10.3389/fpsyg.2014.00261

Guglielmotti, V., & Fiorino, L. (1998). Asymmetry in the Left and Right Habenulo-Interpeduncular Tracts in the Frog. Brain Research Bulletin, 45(1), 105–110. https://doi.org/10.1016/S0361-9230(97)00315-8

He, N., Sethi, S. K., Zhang, C., Li, Y., Chen, Y., Sun, B., Yan, F., & Haacke, E. M. (2020). Visualizing the lateral habenula using susceptibility weighted imaging and quantitative susceptibility mapping. Magnetic Resonance Imaging, 65, 55–61. https://doi.org/10.1016/j.mri.2019.09.005

Hennigan, K., D’Ardenne, K., & McClure, S. M. (2015). Distinct Midbrain and Habenula Pathways Are Involved in Processing Aversive Events in Humans. Journal of Neuroscience, 35(1), 198–208. https://doi.org/10.1523/JNEUROSCI.0927-14.2015

Hétu, S., Luo, Y., Saez, I., D’Ardenne, K., Lohrenz, T., & Montague, P. R. (2016). Asymmetry in functional connectivity of the human habenula revealed by highlJresolution cardiaclJgated resting state imaging. Human Brain Mapping, 37(7), 2602–2615. https://doi.org/10.1002/hbm.23194

Hikosaka, O. (2010). The habenula: From stress evasion to value-based decision-making. Nature Reviews Neuroscience, 11(7), 503–513. https://doi.org/10.1038/nrn2866

Kim, J. H., Lee, J. W., Kim, G. H., Roh, J. H., Kim, M.-J., Seo, S. W., Kim, S. T., Jeon, S., Lee, J.-M., Heilman, K. M., & Na, D. L. (2012). Cortical asymmetries in normal, mild cognitive impairment, and Alzheimer’s disease. Neurobiology of Aging, 33(9), 1959–1966. https://doi.org/10.1016/j.neurobiolaging.2011.06.026

Kim, J., Naidich, T. P., Ely, B. A., Yacoub, E., De Martino, F., Fowkes, M. E., Goodman, W. K., & Xu, J. (2016). Human habenula segmentation using myelin content. NeuroImage, 130, 145–156. https://doi.org/10.1016/j.neuroimage.2016.01.048

Kim, J.-W., Naidich, T. P., Joseph, J., Nair, D., Glasser, M. F., O’halloran, R., Doucet, G. E., Lee, W. H., Krinsky, H., Paulino, A., Glahn, D. C., Anticevic, A., Frangou, S., & Xu, J. (2018). Reproducibility of myelin content-based human habenula segmentation at 3 Tesla. Human Brain Mapping, 39(7), 3058–3071. https://doi.org/10.1002/hbm.24060

Knapp, G., & Hartung, J. (2003). Improved tests for a random effects meta-regression with a single covariate. Statistics in Medicine, 22(17), 2693–2710. https://doi.org/10.1002/sim.1482

Kong, X., ENIGMA Laterality Working Group, Francks, C., Kong, X., Mathias, S. R., Guadalupe, T., Abé, C., Agartz, I., Akudjedu, T. N., Aleman, A., Alhusaini, S., Allen, N. B., Ames, D., Andreassen, O. A., Vasquez, A. A., Armstrong, N. J., Asherson, P., Bergo, F., Bastin, M. E., … Sellgren, C. (2022). Reproducibility in the absence of selective reporting: An illustration from largelJscale brain asymmetry research. Human Brain Mapping, 43(1), 244–254. https://doi.org/10.1002/hbm.25154

Kong, X., Postema, M. C., Guadalupe, T., Kovel, C., Boedhoe, P. S. W., Hoogman, M., Mathias, S. R., Rooij, D., Schijven, D., Glahn, D. C., Medland, S. E., Jahanshad, N., Thomopoulos, S. I., Turner, J. A., Buitelaar, J., Erp, T. G. M., Franke, B., Fisher, S. E., Heuvel, O. A., … Francks, C. (2022). Mapping brain asymmetry in health and disease through the ENIGMA consortium. Human Brain Mapping, 43(1), 167–181. https://doi.org/10.1002/hbm.25033

Kong, X.-Z., Postema, M., Schijven, D., Castillo, A. C., Pepe, A., Crivello, F., Joliot, M., Mazoyer, B., Fisher, S. E., & Francks, C. (2021). Large-Scale Phenomic and Genomic Analysis of Brain Asymmetrical Skew. Cerebral Cortex, 31(9), 4151–4168. https://doi.org/10.1093/cercor/bhab075

Kong, X.-Z., Mathias, S. R., Guadalupe, T., ENIGMA Laterality Working Group, Glahn, D. C., Franke, B., Crivello, F., Tzourio-Mazoyer, N., Fisher, S. E., Thompson, P. M., Francks, C., ENIGMA Laterality Working Group, Kong, X.-Z., Mathias, S. R., Guadalupe, T., Abé, C., Agartz, I., Akudjedu, T. N., Aleman, A., … Orhan, F. (2018). Mapping cortical brain asymmetry in 17,141 healthy individuals worldwide via the ENIGMA Consortium. Proceedings of the National Academy of Sciences, 115(22). https://doi.org/10.1073/pnas.1718418115

Lawson, R. P., Drevets, W. C., & Roiser, J. P. (2013). Defining the habenula in human neuroimaging studies. NeuroImage, 64, 722–727. https://doi.org/10.1016/j.neuroimage.2012.08.076

Lawson, R. P., Nord, C. L., Seymour, B., Thomas, D. L., Dayan, P., Pilling, S., & Roiser, J. P. (2017). Disrupted habenula function in major depression. Molecular Psychiatry, 22(2), 202–208. https://doi.org/10.1038/mp.2016.81

Lawson, R. P., Seymour, B., Loh, E., Lutti, A., Dolan, R. J., Dayan, P., Weiskopf, N., & Roiser, J. . (2014). The habenula encodes negative motivational value associated with primary punishment in humans. Proceedings of the National Academy of Sciences, 111(32), 11858–11863. https://doi.org/10.1073/pnas.1323586111

Leonard, C. M., & Eckert, M. A. (2008). Asymmetry and Dyslexia. Developmental Neuropsychology, 33(6), 663–681. https://doi.org/10.1080/87565640802418597

Lim, S.-H., Yoon, J., Kim, Y. J., Kang, C.-K., Cho, S.-E., Kim, K. G., & Kang, S.-G. (2021). Reproducibility of automated habenula segmentation via deep learning in major depressive disorder and normal controls with 7 Tesla MRI. Scientific Reports, 11(1), 13445. https://doi.org/10.1038/s41598-021-92952-z

Liu, W.-H., Valton, V., Wang, L.-Z., Zhu, Y.-H., & Roiser, J. P. (2017). Association between habenula dysfunction and motivational symptoms in unmedicated major depressive disorder. Social Cognitive and Affective Neuroscience, 12(9), 1520–1533. https://doi.org/10.1093/scan/nsx074

Luan, S., Zhang, L., Wang, R., Zhao, H., & Liu, C. (2019). A restinglJstate study of volumetric and functional connectivity of the habenular nucleus in treatmentlJresistant depression patients. Brain and Behavior, 9(4), e01229. https://doi.org/10.1002/brb3.1229

Menzies, L., Williams, G. B., Chamberlain, S. R., Ooi, C., Fineberg, N., Suckling, J., Sahakian, B. J., Robbins, T. W., & Bullmore, E. T. (2008). White Matter Abnormalities in Patients With Obsessive-Compulsive Disorder and Their First-Degree Relatives. American Journal of Psychiatry, 165(10), 1308–1315. https://doi.org/10.1176/appi.ajp.2008.07101677

Mizumori, S. J. Y., & Baker, P. M. (2017). The Lateral Habenula and Adaptive Behaviors. Trends in Neurosciences, 40(8), 481–493. https://doi.org/10.1016/j.tins.2017.06.001

Müller, U. J., Ahrens, M., Vasilevska, V., Dobrowolny, H., Schiltz, K., Schlaaff, K., Mawrin, C., Frodl, T., Bogerts, B., Gos, T., Truebner, K., Bernstein, H.-G., & Steiner, J. (2021). Reduced habenular volumes and neuron numbers in male heroin addicts: A post-mortem study. European Archives of Psychiatry and Clinical Neuroscience, 271(5), 835–845. https://doi.org/10.1007/s00406-020-01195-y

Ocklenburg, S., Berretz, G., Packheiser, J., & Friedrich, P. (2021). Laterality 2020: Entering the next decade. Laterality, 26(3), 265–297. https://doi.org/10.1080/1357650X.2020.1804396

Ocklenburg, S., & Güntürkün, O. (2018). Sex Differences in Hemispheric Asymmetries. In The Lateralized Brain (pp. 289–311). Elsevier. https://doi.org/10.1016/B978-0-12-803452-1.00011-4

Page, M. J., McKenzie, J. E., Bossuyt, P. M., Boutron, I., Hoffmann, T. C., Mulrow, C. D., Shamseer, L., Tetzlaff, J. M., Akl, E. A., Brennan, S. E., Chou, R., Glanville, J., Grimshaw, J. M., Hróbjartsson, A., Lalu, M. M., Li, T., Loder, E. W., Mayo-Wilson, E., McDonald, S., … Moher, D. (2021). The PRISMA 2020 statement: An updated guideline for reporting systematic reviews. BMJ, n71. https://doi.org/10.1136/bmj.n71

Pipitone, J., Park, M. T. M., Winterburn, J., Lett, T. A., Lerch, J. P., Pruessner, J. C., Lepage, M., Voineskos, A. N., & Chakravarty, M. M. (2014). Multi-atlas segmentation of the whole hippocampus and subfields using multiple automatically generated templates. NeuroImage, 101, 494–512. https://doi.org/10.1016/j.neuroimage.2014.04.054

Ranft, K., Dobrowolny, H., Krell, D., Bielau, H., Bogerts, B., & Bernstein, H.-G. (2010). Evidence for structural abnormalities of the human habenular complex in affective disorders but not in schizophrenia. Psychological Medicine, 40(4), 557–567. https://doi.org/10.1017/S0033291709990821

Sartorius, A., & Henn, F. A. (2007). Deep brain stimulation of the lateral habenula in treatment resistant major depression. Medical Hypotheses, 69(6), 1305–1308. https://doi.org/10.1016/j.mehy.2007.03.021

Sartorius, A., Kiening, K. L., Kirsch, P., von Gall, C. C., Haberkorn, U., Unterberg, A. W., Henn, F. A., & Meyer-Lindenberg, A. (2010). Remission of Major Depression Under Deep Brain Stimulation of the Lateral Habenula in a Therapy-Refractory Patient. Biological Psychiatry, 67(2), e9–e11. https://doi.org/10.1016/j.biopsych.2009.08.027

Savitz, J. B., Bonne, O., Nugent, A. C., Vythilingam, M., Bogers, W., Charney, D. S., & Drevets W. C. (2011a). Habenula volume in post-traumatic stress disorder measured with high-resolution MRI. Biology of Mood & Anxiety Disorders, 1(1), 7. https://doi.org/10.1186/2045-5380-1-7

Savitz, J. B., Nugent, A. C., Bogers, W., Roiser, J. P., Bain, E. E., Neumeister, A., Zarate, C. A., Manji, H. K., Cannon, D. M., Marrett, S., Henn, F., Charney, D. S., & Drevets, W. C. (2011b). Habenula Volume in Bipolar Disorder and Major Depressive Disorder: A High-Resolution Magnetic Resonance Imaging Study. Biological Psychiatry, 69(4), 336–343. https://doi.org/10.1016/j.biopsych.2010.09.027

Schafer, M., Kim, J.-W., Joseph, J., Xu, J., Frangou, S., & Doucet, G. E. (2018). Imaging Habenula Volume in Schizophrenia and Bipolar Disorder. Frontiers in Psychiatry, 9, 456. https://doi.org/10.3389/fpsyt.2018.00456

Schmidt, F. M., Schindler, S., Adamidis, M., Strauß, M., Tränkner, A., Trampel, R., Walter, M., Hegerl, U., Turner, R., Geyer, S., & Schönknecht, P. (2016). Habenula volume increases with disease severity in unmedicated major depressive disorder as revealed by 7T MRI. European Archives of Psychiatry and Clinical Neuroscience, 267(2), 107–115. https://doi.org/10.1007/s00406-016-0675-8

Sudlow, C., Gallacher, J., Allen, N., Beral, V., Burton, P., Danesh, J., Downey, P., Elliott, P., Green, J., Landray, M., Liu, B., Matthews, P., Ong, G., Pell, J., Silman, A., Young, A., Sprosen, T., Peakman, T., & Collins, R. (2015). UK Biobank: An Open Access Resource for Identifying the Causes of a Wide Range of Complex Diseases of Middle and Old Age. PLOS Medicine, 12(3), e1001779. https://doi.org/10.1371/journal.pmed.1001779

Thompson, P. M., Jahanshad, N., Ching, C. R. K., Salminen, L. E., Thomopoulos, S. I., Bright, J., Baune, B. T., Bertolín, S., Bralten, J., Bruin, W. B., Bülow, R., Chen, J., Chye, Y., Dannlowski, U., de Kovel, C. G. F., Donohoe, G., Eyler, L. T., Faraone, S. V., … Zelman, V. (2020). ENIGMA and global neuroscience: A decade of large-scale studies of the brain in health and disease across more than 40 countries. Translational Psychiatry, 10(1), 100. https://doi.org/10.1038/s41398-020-0705-1

Toga, A. W., & Thompson, P. M. (2003). Mapping brain asymmetry. Nature Reviews Neuroscience, 4(1), 37–48. https://doi.org/10.1038/nrn1009

Torrisi, S., Nord, C. L., Balderston, N. L., Roiser, J. P., Grillon, C., & Ernst, M. (2017). Resting state connectivity of the human habenula at ultra-high field. NeuroImage, 147, 872–879. https://doi.org/10.1016/j.neuroimage.2016.10.034

Viechtbauer, W. (2005). Bias and Efficiency of Meta-Analytic Variance Estimators in the Random-Effects Model. Journal of Educational and Behavioral Statistics, 30(3), 261–293. https://doi.org/10.3102/10769986030003261

Wree, A., Zilles, K., & Schleicher, A. (1981). Growth of fresh volumes and spontaneous cell death in the nuclei habenulae of albino rats during ontogenesis. Anatomy and Embryology, 161(4), 419–431. https://doi.org/10.1007/BF00316052

Yoshino, A., Okamoto, Y., Sumiya, Y., Okada, G., Takamura, M., Ichikawa, N., Nakano, T., Shibasaki, C., Aizawa, H., Yamawaki, Y., Kawakami, K., Yokoyama, S., Yoshimoto, J., & Yamawaki, S. (2020). Importance of the Habenula for Avoidance Learning Including Contextual Cues in the Human Brain: A Preliminary fMRI Study. Frontiers in Human Neuroscience, 14, 165. https://doi.org/10.3389/fnhum.2020.00165

Zhang, L., Wang, H., Luan, S., Yang, S., Wang, Z., Wang, J., & Zhao, H. (2017). Altered Volume and Functional Connectivity of the Habenula in Schizophrenia. Frontiers in Human Neuroscience, 11, 636. https://doi.org/10.3389/fnhum.2017.00636

